# Extracellular GABA waves regulate coincidence detection in excitatory circuits

**DOI:** 10.1101/2020.06.15.152652

**Authors:** Sergiy Sylantyev, Leonid P. Savtchenko, Nathanael O’Neill, Dmitri A. Rusakov

## Abstract

Coincidence detection of excitatory inputs by principal neurons underpins the rules of signal integration and Hebbian plasticity in the brain. In the hippocampal circuitry, detection fidelity is thought to depend on the GABAergic synaptic input through a feed-forward inhibitory circuit also involving the hyperpolarization-activated *I*_*h*_ current. However, afferent connections often bypass feed-forward circuitry, suggesting that a different GABAergic mechanism might control coincidence detection in such cases. To test whether fluctuations in the extracellular GABA concentration [GABA] could play a regulatory role here, we use a GABA ‘sniffer’ patch in acute hippocampal slices of the rat and document strong dependence of [GABA] on network activity. We find that blocking GABAergic signalling strongly reduces the coincidence detection window of direct excitatory inputs to pyramidal cells whereas increasing [GABA] through GABA uptake blockade expands it. The underlying mechanism involves membrane-shunting tonic GABA_A_ receptor current; it does not have to rely on *I*_*h*_ but depends strongly on the neuronal GABA transporter GAT-1. We use dendrite-soma dual patch-clamp recordings to show that the strong effect of membrane shunting on coincidence detection relies on nonlinear amplification of changes in the decay of dendritic synaptic currents when they reach the soma. Our results suggest that, by dynamically regulating extracellular GABA, brain network activity can optimise signal integration rules in local excitatory circuits.

## INTRODUCTION

High-precision input coincidence detection by principal neurons is essential for faithful information transfer by brain circuits (Konig *et al*., 1996). The timing and sequence of near-coincident pre- and postsynaptic spikes also controls long-lasting changes of synaptic efficacy (Bi & Poo, 1998). Coincidence detection fidelity, at least in the well-explored hippocampal CA3-CA1 circuit, has been shown to depend on feed-forward inhibition (Pouille & Scanziani, 2001), which is manifested as the biphasic EPSP-IPSPs recorded in the postsynaptic CA1 pyramidal cells (PCs) (Alger & Nicoll, 1982). In the EPSP-IPSP response, the later IPSP component curtails the excitatory conductance decay thus providing a sharper waveform for temporal signal integration. In addition, membrane shunting by the hyperpolarization-activated current *I*_*h*_ (Robinson & Siegelbaum, 2003) accelerates the IPSP component, thus further narrowing the input integration time window (Pavlov *et al*., 2011). The strong influence of shunting conductance on coincidence detection was also demonstrated using dynamic-clamp somatic current injections in cortical PCs (Grande *et al*., 2004), and in electrically compact neurons of the chicken nucleus laminaris (Tang *et al*., 2011).

Intriguingly, the critical role of feed-forward inhibition in coincidence detection fidelity in the CA3-CA1 circuit has been discovered using extracellular stimulation of Schaffer collaterals (Pouille & Scanziani, 2001; Pavlov *et al*., 2011). In contrast, paired CA3-CA1 PC recordings in organotypic hippocampal slices (Debanne *et al*., 1996; Zhang *et al*., 2008), or selective optogenetic stimulation of CA3 PCs or Schaffer collateral axons in acute slices (Kohl *et al*., 2011; Jackman *et al*., 2014) produce robust monophasic EPSPs sufficient for PC spiking (Jackman *et al*., 2014), with no contribution from the intact inhibitory circuitry. Nor do CA1 PCs recorded *in vivo* appear to display biphasic EPSP-IPSP responses as a prevalent feature (Bahner *et al*., 2011; Kowalski *et al*., 2016). These observations indicate that physiological activity of Schaffer collaterals does not necessarily engage feed-forward inhibitory interneurons, instead activating CA1 pyramidal cells directly. The question therefore arises whether local network activity other than feed-forward inhibition, can control coincidence detection of direct, monosynaptic excitatory inputs to principal neurons.

One powerful mechanism that generates sustained membrane-shunting conductance in principal neurons, in particular in hippocampal PCs, is tonic GABA_A_ receptor current (Semyanov *et al*., 2003; Scimemi *et al*., 2005; Glykys & Mody, 2007). This tonic current arises from the incessant bombardment of GABA_A_ receptors by GABA molecules that diffuse from remote synaptic sources, or sometimes released stochastically from local synapses whose individual IPSCs are indistinguishable from noise. In this context, one important feature of GABAergic synapses is that GABA normally escapes the synaptic cleft activating target receptors within at least a several-micron wide volume of tissue (Olah *et al*., 2009). Tonic GABA current thus depends on the extracellular GABA concentration ([GABA]), which reflects the balance of network-driven GABA release and uptake (Glykys & Mody, 2007; Pavlov *et al*., 2014). However, it remains unclear to what degree the local network activity could dynamically control [GABA], given the sparsity of direct inhibitory inputs. A recent study employed a genetically encoded optical GABA sensor to detect a relatively brief (200-300 ms) rise in [GABA] in response to epileptiform discharges in the cortex (Marvin *et al*., 2019). Whether such short transients are indeed characteristic for extracellular GABA waves or whether their detection has been curtailed by the relatively low sensitivity of the sensor remains to be ascertained (Marvin *et al*., 2019).

Whether the [GABA]-dependent tonic membrane current influences the coincidence detection to a significant degree is not a trivial question. Blockade of GABA_A_ receptors alters the holding current in CA1 pyramidal cells by only 5-10 pA (Semyanov *et al*., 2003; Scimemi *et al*., 2005). The expected effect of this change on the time course of local dendritic synaptic currents is likely to be in the sub-millisecond or low millisecond range (Tran-Van-Minh *et al*., 2016). If dendritic signals were to undergo passive filtering while arriving at the soma, this small change would remain such, which would unlikely to affect coincidence detection fidelity. However, blocking a similarly small membrane-shunting influence of *I*_*h*_ changes the coincidence detection window by tens of milliseconds (Pouille & Scanziani, 2001; Pavlov *et al*., 2011), a phenomenon ascribed to active mechanisms of dendritic integration (Magee, 1999; Angelo *et al*., 2007). In this context, it would seem important to understand whether active dendritic filtering is a universal mechanism that amplifies changes in the local synaptic signal time course (such as changes triggered by [GABA] fluctuations), independently of their origin.

To address these issues, first, we implemented a highly sensitive outside-out GABA ‘sniffer’ patch (Isaacson *et al*., 1993; Wlodarczyk *et al*., 2013) to evaluate the extent of activity-dependent extracellular GABA fluctuations in hippocampal tissue. Second, we established the relationship between tonic GABA conductance and the integration time window for direct excitatory inputs to CA1 PCs, and the contributing regulatory role of GABA transporters. Finally, we examined Schaffer collateral-elicited EPSCs in dual dendrite-soma patch recordings of CA1 PCs, to explore amplification of small kinetic changes in local synaptic currents in the course of dendritic integration.

## METHODS

### Animals and electrophysiology

Animal procedures were conducted in accordance with the United Kingdom Home Office (Scientific Procedures) Act (1986) Schedule 1, in full compliance with the relevant ethics policiesUCL and the University of Edinburgh ethical committee regulations. 3-4-week old male Sprague-Dawley or Wistar rats were bred in the institutional animal house, grown on a Rat and Mouse Breeding Diet (Special Diet Services, Witham, UK) and water ad libitum, and maintained at 12–12-h light-dark (L/D) cycle. Animals were sacrificed in the first half of light period of the L/D cycle with an overdose of isoflurane. After decapitation with guillotine, brains were rapidly removed and dissected, and hippocampi sliced.

Transverse 300 μm hippocampal slices were cut incubated for one hour in a solution containing (in mM): 124 NaCl, 3 KCl, 1 CaCl_2_, 3 MgCl_2_, 26 NaHCO_3_, 1.25 NaH_2_PO_4_, 10 D-glucose, and bubbled with 95:5 O_2_/CO_2_, pH 7.4. Slices were next transferred to a recording chamber superfused with an external solution which was similar to the incubation solution except 2 mM CaCl_2_ and 2 mM MgCl_2_. Where specified, GABA_A_ receptors were blocked with 50 μM picrotoxin (PTX), *I*_*h*_ with 10 μM ZD-7288, and AMPA receptor desensitisation with 10 μM cyclothiazide (CTZ).

The intracellular solution for voltage-clamp recordings contained (mM): 117.5 Cs-gluconate, 17.5 CsCl, 10 KOH-HEPES, 10 BAPTA, 8 NaCl, 5 QX-314 Cl^-^, 2 Mg-ATP, 0.3 GTP (pH 7.2, 295 mOsm); the intracellular solution for current-clamp recordings contained (mM): 126 K-gluconate, 4 NaCl, 5 KOH-HEPES, 10 glucose, 1 MgSO4□7H2O, 2 BAPTA, 2Mg -ATP. Morphological tracer Alexa Fluor 594 was added in some experiments for cell visualisation. Patch-clamp recordings were performed using Multiclamp-700B amplifier; signals were digitized at 10 kHz. The pipette resistance was 3-6 MΩ for whole-cell recordings and 7-9 MΩ for outside-out patches.

Apical dendrites of CA1 pyramidal cells were patched whole-cell 50-150 μm from the soma. Two theta-glass pipette electrodes pulled to 20-40 μm filled with ACSF were used to stimulate Schaffer collaterals with 50-150 μs electrical stimuli; individual recording sweeps were collected at 15 s intervals. Simulation strength was adjusted so that (a) each of the two afferent stimuli produced somatic EPSPs featuring approximately similar amplitudes, and (b) upon coincident stimulation of the two inputs the postsynaptic cell generated an action potential with the probability of >0.9 (which was tested by recording ∼50 trials). In the coincidence-window experiments, 10 trials were routinely recorded for each time interval between the afferent input onsets. Data were represented as mean ± SEM; Student’s unpaired or paired t-test (or non-parametric Wilcoxon paired tests when distribution was non-Gaussian) was used for statistical hypothesis testing.

### Monitoring extracellular GABA with an outside-out ‘sniffer patch’

Outside-out patches were pulled from dentate granule cells, lifted above the slice tissue and carefully lowered into the slice region of interest near the surface, as detailed previously (Sylantyev & Rusakov, 2013; Wlodarczyk et al., 2013). These recordings were performed in voltage-clamp mode at V_h_ = −70 mV. Where specified, GABA_A_R-mediated single-channel currents were recorded in the presence of 0.1 μM CGP-55845, 200 μM S-MCPG and 1 μM strychnine. Single-channel recordings were acquired at 10 kHz, noise >1 kHz was subsequently filtered out off-line. GABA_A_ receptor specificity was routinely confirmed by adding 50 μM PTX at the end of recording.

To assess changes in extracellular GABA, we used the open-time fraction of single channel openings. This was calculated as t_o_/t_f_ ratio, where t_o_ is the overall duration of all individual channel openings over recording time t_f_. Some patches contained more than one GABA_A_R channel and could therefore display simultaneous multiple channel openings: in such cases, all individual channel-opening durations were still added up, thus allowing the t_o_/t_f_ value to exceed one. Single-channel openings were selected automatically by the threshold-detection algorithm of Clampfit software (Molecular Devices), with the minimum event duration of 0.2 ms, and the channel-opening current threshold set at 1.5 pA. In one set of experiments, as indicated, we used a ‘large’ sniffer patch to boost the number of GABA_A_ receptors: this involved obtaining a nucleated outside-out patch from dentate gyrus granule cell in the same slice, as described before (Sun *et al*., 2020).

The burst protocol included eight trains of 10 pulses at 100 Hz, 1 s apart, delivered by a bipolar electrode placed in the stratum radiatum; this stimulation was below the threshold necessary for the induction of long-term synaptic potentiation. The average GABA_A_ receptor-channel open time fraction was calculated over a five-second interval after the eighth burst. To prompt spontaneous network discharges, we perfused slices with a Mg-free aCSF containing 5 mM KCl; single-channel openings were recorded before the development of any epileptiform activity in the slice. All recordings were done at 32-33°C. Field potential recordings from CA1 stratum pyramidale were performed with 1-2 MΩ glass electrodes filled with aCSF. ZD-7288, CTZ, SNAP-5114, SKF-89976A, NBQX, DNQX, CGP-55845, S-MCPG, strychnine and PTX were purchased from Tocris Bioscience. All other chemicals were purchased from Sigma-Aldrich.

### NEURON modelling: pyramidal cell

Simulations were performed on a 3D-reconstructed pyramidal neuron from the hippocampal area CA1 (https://senselab.med.yale.edu/modeldb, NEURON accession number 7509) (Magee & Cook, 2000), with excitatory synapses distributed over the dendritic tree. The cell axial specific resistance and capacitive were, respectively, *R*_*a*_ *=* 90 Om cm, and *C*_*m*_ = 1 µF/cm^2^. Excitatory synaptic conductance time course *g*_*s*_*(t)* was modelled using the NEURON 7.0 function Exp2Syn (dual-exponential): *gs* (*t*) = *Gs* (exp(−*t* / *τ*1) − exp(−*t* / *τ*2)) where *τ*_*1*_ and *τ*_*2*_ are the rise and decay time constants, respectively, and *G*_*s*_ is the synaptic combined conductance. The values of *τ*_*1*_ and *τ*_*2*_ were established empirically, by matching simulated somatic and dendritic EPSPs with experimental recordings (note that synaptic conductance does not necessarily follow AMPA receptor kinetics in response to sub millisecond glutamate pulses in outside-out patches (Sylantyev *et al*., 2008; Sylantyev *et al*., 2013)). This procedure led to setting the values of *τ*_*1*_ and *τ*_*2*_ at 2.5 ms and 10 ms, respectively. The reverse potential of excitatory synapses was set at 0.

### Modelling tonic GABA_A_ and I_h_ receptor currents

Tonic GABA_A_ receptor-mediated current (reversal potential *E*_GABA_ is between -75 and −55 mV) is an outwardly rectifying shunting current routinely detected in principal neurons (Semyanov *et al*., 2003; Sylantyev *et al*., 2008; Pavlov *et al*., 2009; Sylantyev *et al*., 2013). Its conductance *I*_GABA_ = *g*_GABA_× *O* × (*V* − *E*_GABA_) was calculated using *g*_GABA_ = 3 mS cm^-2^, where the state *O* is a proportion of channels in the open state, as estimated previously in CA1 pyramidal cells (Pavlov *et al*., 2009; Song *et al*., 2011). The transition from the open to the close state was described by a straightforward kinetic scheme reported earlier (Pavlov *et al*., 2009).

The kinetic of *I*_*h*_ (the hyperpolarization-activated cation current) was copied from the 3D-reconstructed NEURON cell model (accession number, 7509), which was optimised to fit experimental observations (Magee, 1999). The deactivation potential of *I*_*h*_ was -81 mV, unit conductance 0.1 mS cm^-2^, in line with previously published estimates (Magee, 1999; Pavlov *et al*., 2011).

### Simulating coincidence detection

The simulated pyramidal cell was equipped with 40 excitatory synapses scattered along apical dendrites and divided into two separate equal groups, 20 synapses each, to mimic two independent afferent inputs. A random number generator was used to activate synaptic inputs with the average probability P_r_ = 0.35. The synaptic conductance of individual synapses was adjusted to induce a postsynaptic spike with >0.9 probability upon the coincident activation of the two synaptic groups. In practice, we tested this using around ∼100 trials achieving the spike success rate between 95 and 99. Routinely, one synaptic group was activated at 50 ms after the ‘sweep’ onset, with the other group activate at different time points, between 30 ms before and 30 ms after the first group onset. To estimate the coincidence detection window, the spike probability was calculated, throughout varied time intervals, 10 times (which thus produced 10 different sets of stochastically activated synapses, on average 0.35.20 = 7 synapses per trial), which was similar to the experimental design used.

### Statistical summary and software accessibility

Experiments involved straightforward statistical (paired-sample) designs, with the statistical units represented by individual cells (one cell per acute slice), which contribute the main source of variance with respect to the variable of interest. Sampling was quasi-random, in line with the established criteria for patch-clamp experiments. Leaky cells (holding current >20-30 pA at V_h_ = -65 mV) were discarded. The data were routinely presented as a scatter of measurements from individual cells or patches, and additionally described as mean ± SEM. The width of coincidence windows was represented by the best-fit Gaussian dispersion σ, and the average window profile was displayed as mean ± SEM (number of cells) at each time point whereas the average σ estimate was shown as mean ± SD. To compare statistically the coincidence windows between experimental epochs, we calculated σ value in each individual cell (as the underlying window shape model), which thus represented a statistical unit. Statistical hypotheses pertinent to the mean difference involved a paired-sample t-test or one-way ANOVA, as indicated (OriginPro, OriginLab RRID: SCR_014212). All software codes are available on request and will be deposited for free access upon publication.

## RESULTS

### Neuronal activity can elevate extracellular GABA level several-fold

To understand the magnitude of activity-dependent fluctuations in [GABA], we used a highly sensitive GABA ‘sniffer’, an outside-out cell membrane patch held by the recording micro-pipette in the extracellular medium (Isaacson *et al*., 1993; Wlodarczyk *et al*., 2013) (Fig. 1A). The sniffer patch reports the opening of individual GABA_A_ receptor channels, with the event frequency varying with [GABA]. With a consistent procedure of pipette preparation and patch pulling (Sylantyev & Rusakov, 2013), its [GABA] sensitivity can be calibrated (Fig. 1B). The sniffer-patch method thus reports the dynamics of volume-average [GABA] in the extracellular space adjacent to the patch.

**Figure 1.**
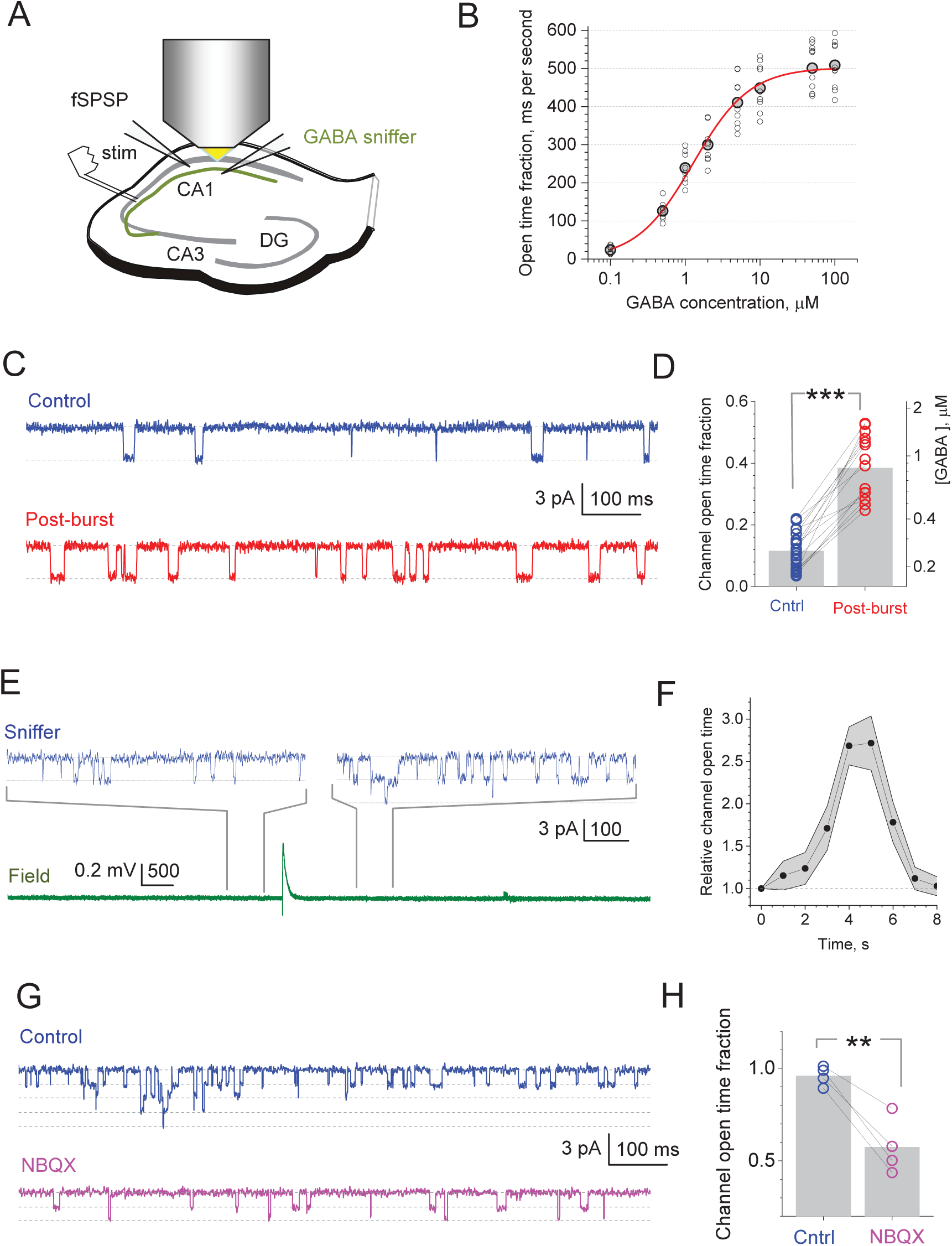
GABA sniffer detects several-fold fluctuations in the extracellular GABA level induced by neural activity changes. **A**, Experiment diagram illustrating recordings in an acute hippocampal slice, with the ‘sniffer patch’ (Methods) held in the extracellular space. **B**, Calibration of the GABA ‘sniffer’ patch: average values of the open time fraction (grey circles, mean; smaller hollow circles, individual data; n = 10 cells), expressed in millisecond per second; red line, best-fit Hill approximation; small variability points to a highly reproducible sniffer-patch protocol. **C**, Typical single-channel activity (1 s interval shown) recorded in experiments as in **a**, inC baseline conditions (Control) and within 5 seconds after electrical stimulation of Schaffer collaterals (post-burst; 8 series of 10 pulses at 100 Hz, 1 s apart). Dotted lines, GABAR channel closed and open current levels. **D**, Summary of experiments shown in **c**: average channel open-time fraction over the 5 s interval post-burst; grey bars, mean values; straight lines connect same-patch experiments; ***p < 0.001 (n = 27 patches in control, including n = 15 paired control / post-burst patches). **E**, Upper traces illustrate sniffer patch recordings sampled before and after a single spontaneous synchronous network discharge shown in the bottom trace (field potential recorded simultaneously, Mg-free bath solution, Methods); sampling time windows are indicated by grey connecting lines. **F**, Time course of the GABA_A_R channel opening kinetics (mean ± SEM, n = 6 cells) after the network discharge as shown in **E** (onset at *t* = 0). **G**, Typical single-channel activity (1 s interval shown) recorded with a sniffer patch that is larger than that in A-D (thus, calibration in B does not apply), in baseline conditions (Control) and after adding 10 μM NBQX. **H**, Summary of experiments shown in **f**; other notations as in **b**); **p < 0.01 (n = 4). Note that the accumulated ‘channel open-time fraction’ for multiple channels in the large patch could exceed 1.

In the first sniffer-patch experiment, we applied a short series of high frequency stimuli to Schaffer collaterals (eight trains of 10 pulses at 100 Hz, 1 s apart). This increased the GABA receptor open-time fraction in the patch three-fold (mean ± SEM: from 0.12 ± 0.01 to 0.39 ± 0.03), which corresponds to the [GABA] increase from ∼300 nM to ∼900 nM (Fig. 1C-D), for at least five seconds post-burst. A qualitatively similar increase could be routinely observed after a single spontaneous network discharge when we perfused the slice with the Mg-free ACSF to boost its excitability (Fig. 1E-F). These experiments detected [GABA] transients that were an order of magnitude longer than those revealed with the optical GABA sensor in similar conditions (Marvin *et al*., 2019), arguing for the high sensitivity of the present method. In another experiment, we used a much larger sniffer patch (nucleated patch from granule cells; Methods), to boost the baseline GABA_A_ receptor channel-opening rate, and found that the blockade of AMPA receptors with NBQX reduced this rate by half (Fig. 1F-G).

Our results thus argue that boosting neuronal activity can generate 2-3-fold transient changes in tissue-average [GABA], lasting for seconds after the activity boost ends. While demonstrating activity-associated increases in [GABA], these experimental protocols generate variable effects on the scale of seconds, which is not suitable for the millisecond-range monitoring of coincidence detection (Pouille & Scanziani, 2001; Pavlov *et al*., 2011). However, the observed range of [GABA] change (Fig. 1) appears compatible with the effect of blocking GABA transport, which roughly doubles tonic GABA_A_ receptor - mediated current in CA1 PCs, and with the effect of blocking GABA_A_ receptors which removes the current (Semyanov *et al*., 2003; Scimemi *et al*., 2005). Thus, to achieve comparable with our observations (Fig. 1) yet stable changes in [GABA] in both directions, our tests of coincidence detection employed the blockade of either GABA_A_ receptors or GABA transporters, as outlined below.

### Tonic GABA current affects coincidence detection beyond the effect of *I*_*h*_

It has previously been shown that *I*_*h*_ plays a major role in narrowing the coincidence detection window in the CA3-CA1 feed-forward inhibition circuit (Pavlov *et al*., 2011). We asked whether tonic GABA conductance has a similar effect, and if so whether the effects of *I*_*h*_ and GABA occlude. First, to rule out the di-synaptic feedforward inhibition circuit, we used theta-glass bipolar stimulating electrodes (Methods) that provide highly localised engagement of afferent fibres in *s. radiatum*, adjusting their positions so that no IPSP component could be detected throughout (Fig. 2B, diagram and traces). This was in striking contrast with the biphasic EPSP-IPSP responses characteristic for experiments that involve feedforward inhibition (Pouille & Scanziani, 2001; Pavlov *et al*., 2011) (see next section for further control of monosynaptic transmission). Next, we confirmed that the decay of monosynaptic EPSPs in CA1 PCs decelerated under *I*_*h*_ blockade by ZD7288 (ZD) (Magee, 1999), but also found that the GABA_A_ receptor blocker PTX prompted further EPSP deceleration (Fig. 2A). Both effects could be readily replicated in a 3D-reconstructed, realistic NEURON model of a CA1 PC (Magee & Cook, 2000) (https://senselab.med.yale.edu/modeldb, NEURON accession number 7509) (Fig. 2A). These observations suggested that the effect of *I*_*h*_ blockade on the EPSP kinetics does not occlude the effect of tonic GABA_A_ current.

**Figure 2.**
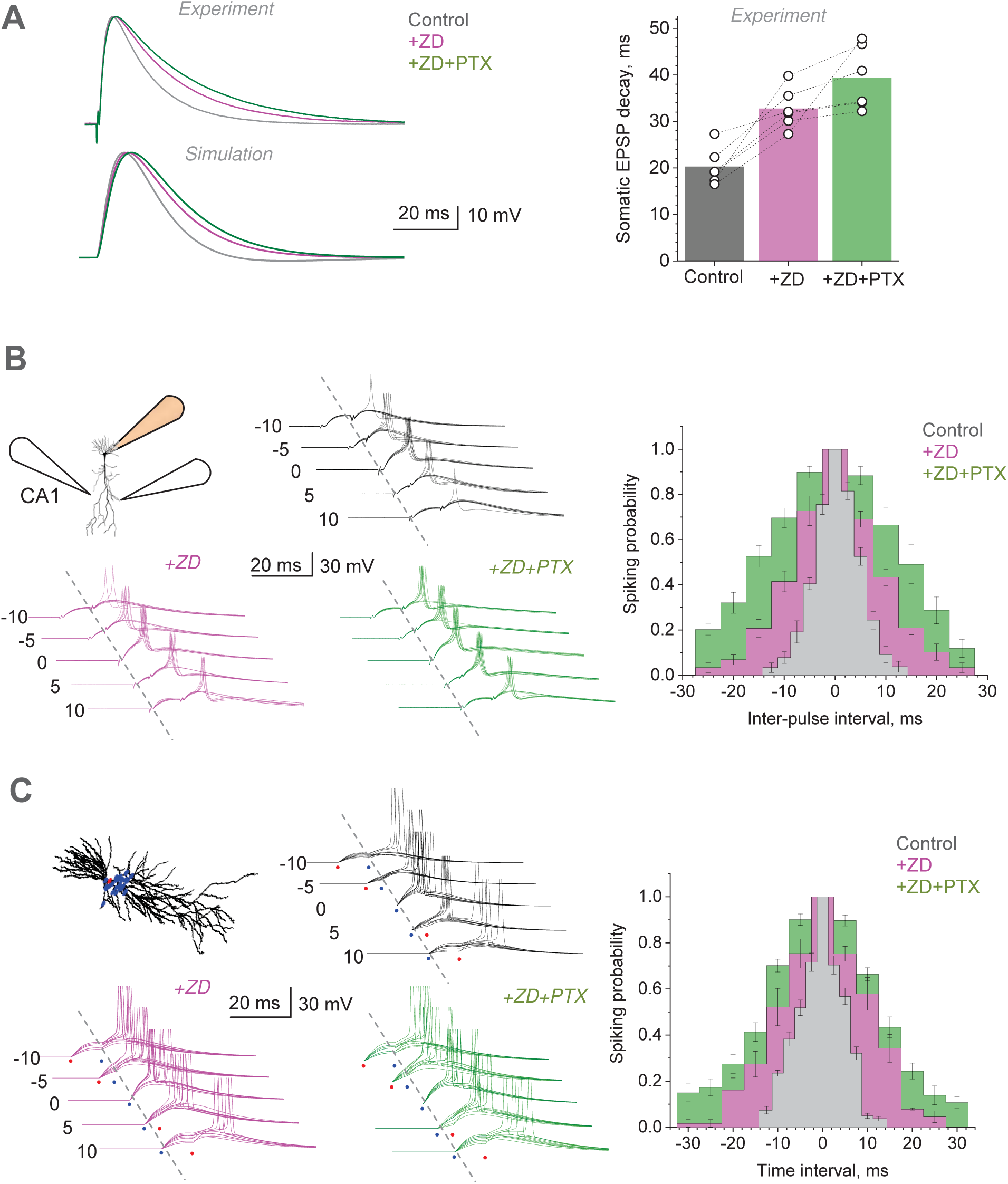
Precision of coincidence detection for distinct excitatory inputs to CA1 pyramidal cells depends on GABA_A_ current. **A**, *Upper traces*: Characteristic EPSPs recorded in the CA1 pyramidal cell soma (upper traces, 10 trial average) in control conditions, after adding 10 μM ZD7288 (+ZD) and subsequently 50 μM PTX (+ZD+PTX), as indicated, normalised to baseline response (scale bar). *Graph*: summary of these experiments (mean and individual data points, n = 6 cells). *Lower traces*: similar tests replicated *in silico* with a NEURON CA1 pyramidal cell model (ModelDB https://senselab.med.yale.edu, accession numbers 2796 and 7509), with 40 synapses scattered along apical dendrites; baseline *I*_*h*_ unit conductance and unit tonic GABA_A_ current are, respectively, 0.1 mS cm^-2^ and ∼3 mS cm^-2^, as estimated earlier ^21, 41, 42^ for quiescent network conditions. **B**, Diagram, experimental design: electrical stimulation of two Schaffer collateral inputs converging onto a CA1 pyramidal cell held in current clamp. Traces, one-cell example of somatic EPSPs (10 consecutive traces), with or without action potentials, at different time intervals between two presynaptic inputs, as indicated (ms), in control conditions (top), after adding 10 μM ZD7288 (+ZD, magenta) and subsequently 50 μM PTX (+ZD+PTX, green), as indicated. Bar graphs, summary of the average spiking probability over the inter-pulse interval (mean ± SEM; n = 6 cells; colour coding as indicated; error bar position shows fixed time points); Gaussian-model paired-sample t-tests show mean σ difference at p < 0.001 for control vs ZD, ZD vs PTX samples (n = 6), and one-way ANOVA for the factor of drug application. **C**, Diagram, simulated CA1 pyramidal cell (NEURON ModelDB https://senselab.med.yale.edu, accession numbers 2796 and 7509) with excitatory inputs (blue dots) scattered across the dendritic tree; excitatory synaptic inputs; conductance time course *Gs* (exp(−*t* / *η*1) − exp(−*t* / *η* 2)) where *τ*_*1*_ = 2.5 ms and *τ*_*2*_ = 10 ms, respectively; *G*_*s*_ is maximal synaptic conductance; release probability P_r_ = 0.35. Traces, simulated EPSPs replicating experiments shown in B, with stochastic synaptic release (10 traces shown for each condition; notations as in B). Bar graphs, the outcome of simulation experiments; coincidence windows for control, +ZD, and +ZD+PTX cases were, respectively: 12.5 ± 0.72, 23.3 ± 0.78, and 29.9 ± 1.15 ms (Gaussian-fit σ ± parameter SD); other notations as in B.

To see how these mechanisms influence coincidence detection of monosynaptic inputs, we sought to stimulate two sets of direct Schaffer collateral connections to CA1 PCs in *s. radiatum* using a similar arrangement for two stimulating theta-glass bipolar electrodes. The stimulus strength was further adjusted to induce a postsynaptic spike with >0.9 probability upon the exact temporal coincidence of the two inputs. As expected, increasing the time interval between the two inputs produced postsynaptic spikes with the progressively decreasing probability. Fitting the spiking probability profile (versus inter-stimulus interval) with the Gaussian gave a temporal coincidence window of 10.1 ± 0.35 ms (Gaussian dispersion σ ± SE; Fig. 2B, grey bars; n = 6 cells). The blockade of *I*_*h*_ with ZD (10 µM) widened the coincidence window to σ = 19.8 ± 0.79 ms (Fig. 2B, magenta bars). A subsequent blockade of GABA_A_ conductance with PTX (50 µM) widened it further, to σ = 31.5 ± 0.87 ms (Fig. 2B, green bars). The latter was similar to the coincidence window reported earlier in the tests under GABAergic transmission blockade (Pouille & Scanziani, 2001).

Again, to see whether these observations are consistent with the known biophysical characteristics of CA1 PCs, we replicated our experiments *in silico*. The modelled reconstructed cell (Magee & Cook, 2000) (see above) was equipped with two independently activated pools of excitatory synapses scattered on its apical dendrites (Fig. 2C, inset). Each synaptic pool contained 20 synapses that could be activated synchronously at a given time. Individual synaptic inputs produced EPSPs stochastically, with a ‘release probability’ of P_r_ = 0.35 (Methods) as estimated earlier (Rusakov & Fine, 2003). These simulations and their outcome faithfully replicated our experiments (Fig. 2C), confirming the biophysical underpinning of our interpretation.

### Tonic GABA current regulates coincidence detection without engaging *I*_*h*_

In the next experiment, we sought to determine whether the tonic GABA current can significantly affect the coincidence detection window when *I*_*h*_ remains intact. Therefore, we added PTX following a recording session in baseline conditions. In such tests, PTX-application increased the EPSP amplitude by only ∼5% while hyperpolarising the cell membrane by 3.52 ± 1.23 mV (Fig. 3A-B), consistent with previous observations (Semyanov *et al*., 2003; Scimemi *et al*., 2005; Pavlov *et al*., 2009). This was in striking contrast with the properties of the biphasic EPSP-IPSP responses generated by the CA3-CA1 excitation and feedforward-inhibition circuit, in which PTX application increases the EPSP amplitude at least two-fold, with a prominent prolongation of the rise time (Pouille & Scanziani, 2001; Pavlov *et al*., 2011). Together with the absence of the IPSC component (Fig. 2A), these data effectively rule out the di-synaptic feedforward inhibition circuitry from our tests. In addition, blocking AMPA and NMDA receptors in our experiments left no evoked signal in CA1 PCs (Fig. 3C), confirming no direct stimulation of local interneurons.

**Figure 3.**
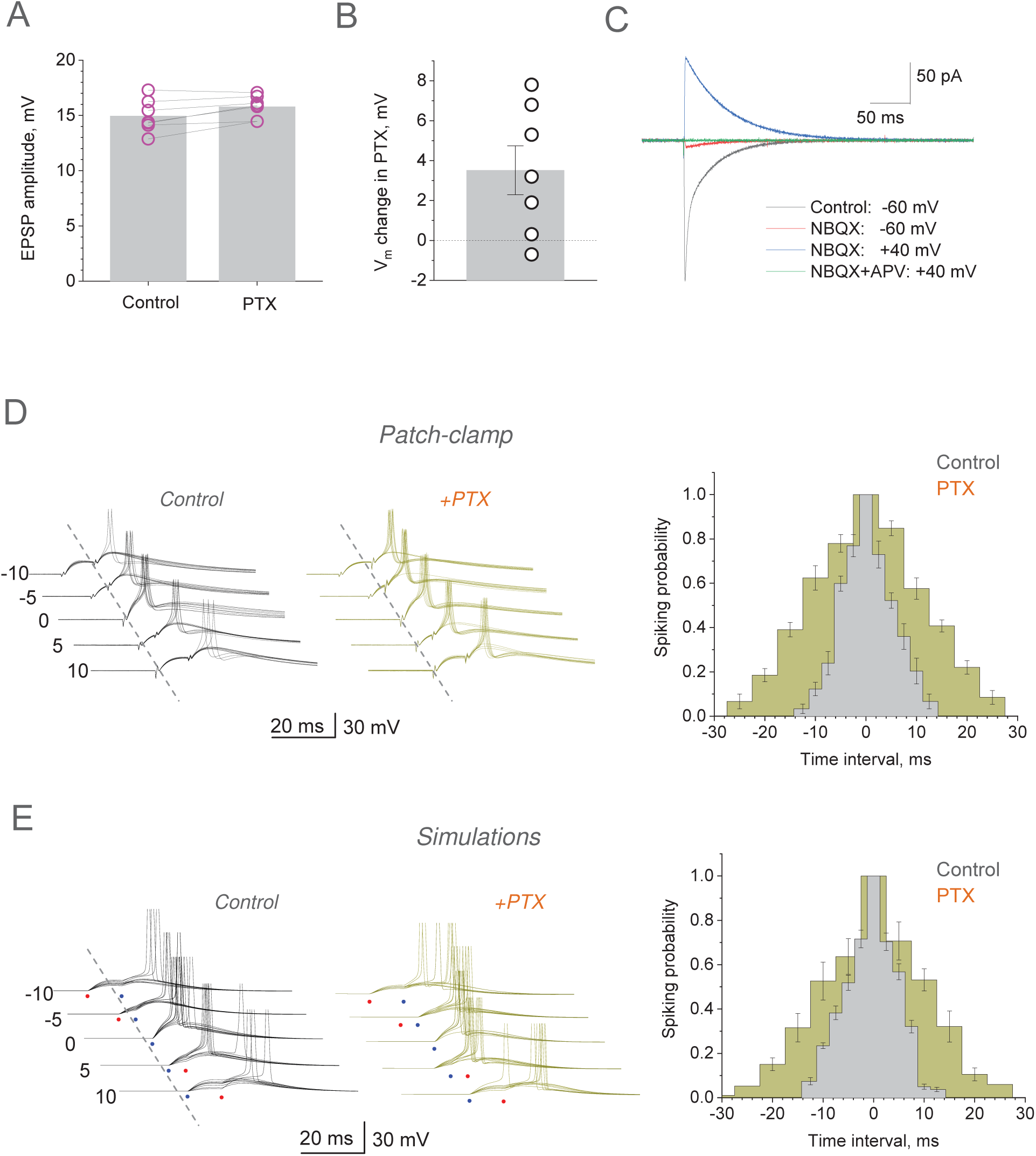
Precision coincidence detection of excitatory inputs by CA1 pyramidal cells depends on tonic GABA_A_ current. **A**, EPSP amplitude in control conditions and after application of 50 μM PTX (mean ± SEM): 15.0 ± 0.56 mV and 15.8 ± 0.38 mV, respectively (n = 7 cells); dots, individual cell data; bars, mean value. **B**, Change in the CA1 pyramidal cell membrane potential V_m_ upon application of 50 μM PTX (mean ± SEM): 3.52 ± 1.23 mV (n = 7). **C, A** test to rule out direct electric stimulation of interneurons (GABA receptors intact, voltage-clamp mode); traces, characteristic EPSCs in control conditions, after application of the AMPA receptor blocker NBQX (20 μM) and subsequent addition of the NMDA receptor blocker APV (50 μM), as indicated. **D**, Traces, one-cell example of somatic EPSPs (10 consecutive traces), with or without generated action potentials, at different time intervals between two presynaptic inputs, as indicated (ms), in control conditions (top) and after adding 50 μM PTX (PTX, green), as indicated. Bar graphs, summary of the average spiking probability over the inter-pulse interval (mean ± SEM; n = 6 cells; colour coding as indicated; error bar position shows fixed time points); Gaussian-model paired-sample t-test shows mean σ difference at p < 0.001 for control vs PTX samples. **E**, Traces, simulated EPSPs replicating experiments in **a**, with stochastic synaptic release (10 traces shown for each condition; notations as in **a**). Histograms, the outcome of simulation experiments; coincidence windows for control and +PTX cases were, respectively: 12.5 ± 0.72 and 24.2 ± 1.2 ms (Gaussian-fit σ ± parameter SD); other notations as in D.

Thus, we found that application of PTX dramatically widened the coincidence window, (σ = 26.7 ± 0.73 ms compared to 12.1 ± 0.55 ms in control; n = 6 cells; Fig. 3D). Because the effect was compatible with that under both *I*_*h*_ and GABA receptor blockade (Fig. 2B), these observations argue that the influence of GABA tonic on coincidence detection does not require *I*_*h*_. The effects of PTX in baseline conditions and under *I*_*h*_ blockade were comparable, arguing for the independent, additive nature of the two mechanisms. Again, simulating these experiments with a realistic CA1 PC model confirmed biophysical underpinning of the observed phenomena (Fig. 3E).

### Coincidence detection is controlled mainly by neuronal GABA transporters

Tonic GABA current depends on [GABA] which is in turn controlled by several types of GABA transporters expressed by nerve and glial cells (Scimemi, 2014) as blocking GABA transport roughly doubles this current in CA1 PCs (Semyanov *et al*., 2003; Scimemi *et al*., 2005). This effect is comparable with the 2-3-fold increase in [GABA] after a burst of network activity (Fig. 1D). Here, we asked therefore whether elevating extracellular GABA by inhibiting GABA transporters would alter the coincidence detection window in our experiments.

Our pilot simulations with the model circuit (as in Figs. 2C and 3E) predicted that increasing membrane shunt from the baseline level could sharply narrow the input coincidence window over which postsynaptic spikes are generated. Ultimately, shortcutting membrane conductance could prevent the postsynaptic cell from firing. Therefore, to avoid a collapse (null-width) of the coincidence window upon the increased shunt, in these experiments we adjusted stimulus strength to start with a relatively wide coincidence interval in baseline conditions. In the first test, we used nipecotic acid (NipA), a GABA transporter blocker, which can also activate GABA_A_ receptors as a false neurotransmitter (Roepstorff & Lambert, 1992). Application of NipA sharply reduced the coincidence window, from σ = 54.8 ± 8.3 to 36.1 ± 4.1 ms (n = 12) whereas the subsequent blockade of GABA_A_ receptors by PTX reversed the effect in the opposite direction, widening the coincidence window to σ = 180.4 ± 12.4 ms (n = 6; the remaining cells were unstable), much beyond that in baseline conditions (Fig. 4A). In the second experiment, blocking the predominantly neuronal GABA transporter GAT-1 with SKF-89976A (SKF) in baseline conditions produced a qualitatively similar effect (changing σ from 123.2 ± 8.3 to 32.9 ± 3.7 ms, n = 9; Fig. 4B). Finally, we asked if glial transporters GAT-3, which have been implicated in controlling extracellular GABA levels under intense network activity (Boddum *et al*., 2016), contribute substantially to the regulation of coincidence detection in baseline conditions. The specific GAT-3 blocker SNAP-5114 did narrow the coincidence widow, from σ = 124.3 ± 6.3 ms to 99.8 ± 5.6 ms (n = 21, Fig. 4C), but it represented only a small fraction of the effect seen with SKF or NipA (Fig. 4A-B). The latter result suggested that neuronal GABA transporters are the main contributor to the regulatory effect of tonic GABA current on the input coincidence window.

**Figure 4.**
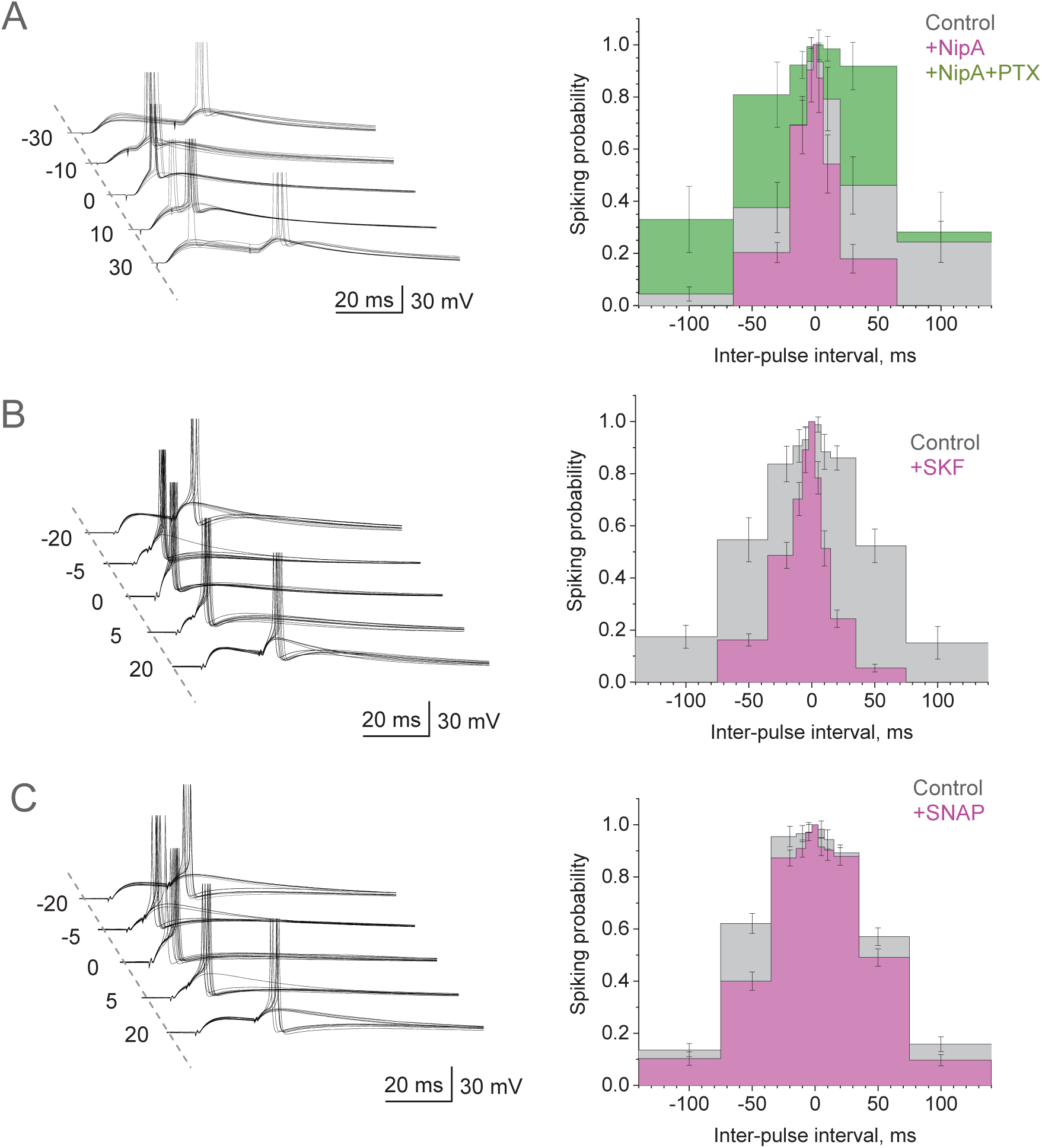
Coincidence detection of excitatory inputs is controlled by GABA transporters. **A**, Traces, characteristic EPSPs (10 consecutive traces), with or without generated spikes, at different time intervals between two stimuli, as indicated (ms), in control conditions. Bar graphs, summary of the average spiking probability over the inter-pulse interval (mean ± SEM) in control conditions (n = 12 cells), with 600 μM nipecotic acid (+NipA; n = 8), and subsequently 50 μM PTX (+NipA+PTX; n = 7); colour coding as indicated; error bar position shows fixed time points.); Gaussian-model paired-sample t-tests showed mean σ difference at p < 0.001 for control vs NipA, p < 0.003 for NipA+PTX vs NipA, and p < 0.001 for one-way ANOVA with the factor of drug application. **B**, Experiment as in **A**, but with 100 μM SKF-89976A (+SKF; n = 9 cells) added. Other notations as in **A**; Gaussian-model paired-sample t-test shows mean σ difference at p < 0.002 for control vs SKF samples. **C**, Experiment as in **A**, but with 100 μM SNAP-5114 (+SNAP; n = 21 cells) added. Other notations as in **A-B;** Gaussian-model paired-sample t-tests showed difference for mean σ at p < 0.001 for control vs SNAP.

### Dendritic processing amplifies small changes in the EPSC decay

Our data indicate that blocking tonic GABA current leads only to a 3-4 mV change in membrane potential (Fig. 3B), consistent with earlier data ascribing 5-10 pA to whole-cell tonic GABA current (Semyanov *et al*., 2003; Scimemi *et al*., 2005). Such a small change is highly unlikely to alter the decay of fast dendritic EPSPs at individual synapses by more than a millisecond (Tran-Van-Minh *et al*., 2016; Jayant *et al*., 2017) yet it prolongs the decay of somatic EPSPs by 5-10 ms (Fig. 2A), which is paralleled by a 10-20 ms change in the coincidence detection window (Figs 2B and 3D). Whether such a small change is indeed amplified when reaching the soma has been a subject of debate: passive filtering does not amplify signal fluctuations. Therefore, to understand whether active filtering is an inherent feature of dendritic integration that can boost small changes in the kinetics of local synaptic currents, regardless of *I*_*h*_ or GABA influence, we carried out dual dendrite-soma whole-cell recordings in CA1 PCs. In these tests, *I*_*h*_ was inhibited by holding cells in voltage-clamp with QX-314 inside (Perkins & Wong, 1995), and GABA_A_ receptors were blocked with 50 µM PTX. Again, we stimulated a single axo-dendritic Schaffer collateral synapse using an extracellular bipolar theta-glass pipette electrode placed within a few microns of the patched and visualised dendrite (Fig. 5A, image). The single-synapse origin of recorded unitary dendritic EPSCs (uEPSCs, voltage-clamp mode) was confirmed by documenting their uniform shape over multiple trials, with a release failure rate of ∼60-70% characteristic for this circuitry (Rusakov & Fine, 2003) (Fig. 5A, traces).

**Figure 5.**
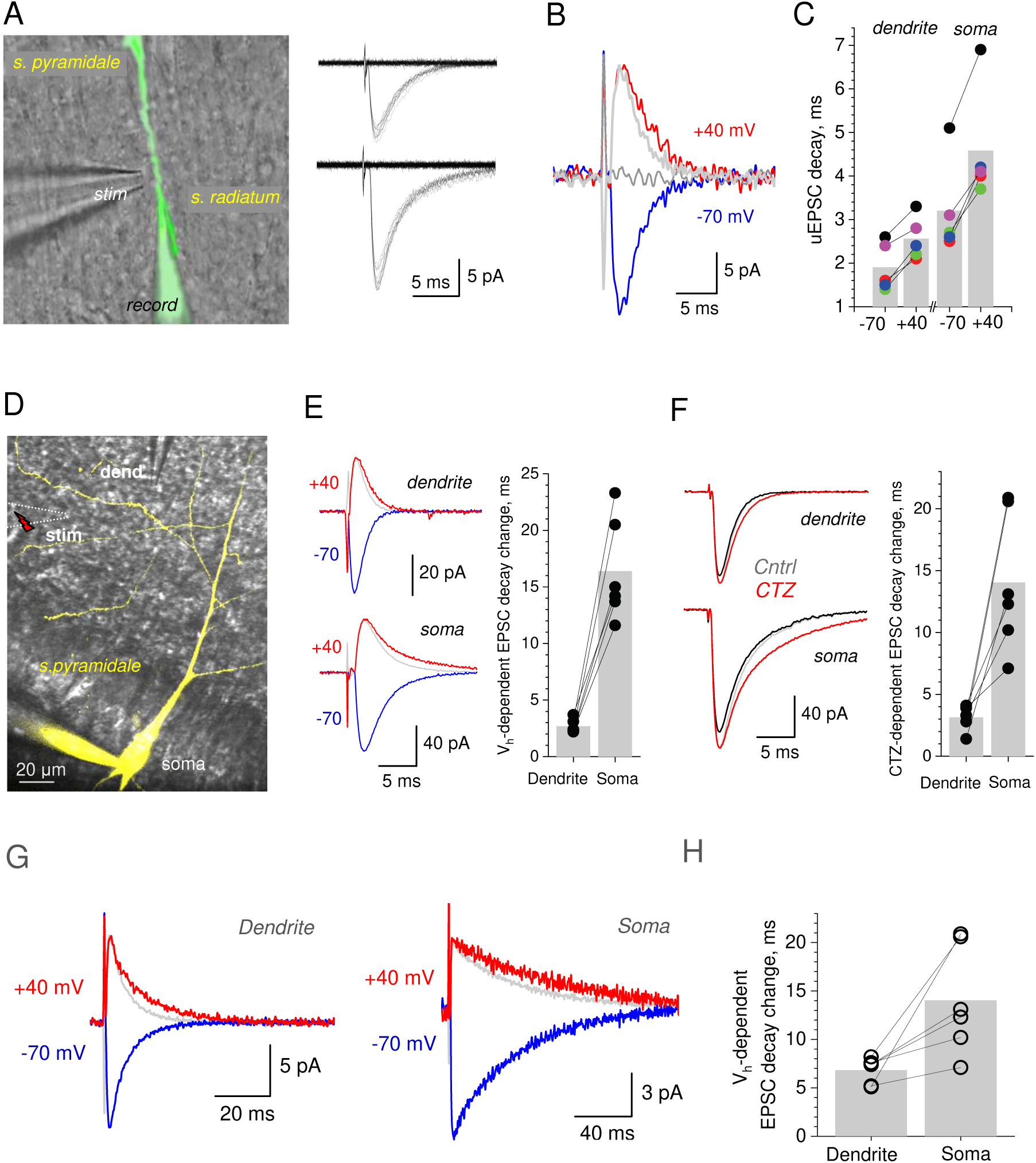
Changes in the dendritic EPSC kinetics are amplified at the soma. **A**, Dendritic-patch recordings of single-synapse, unitary EPSCs (uEPSCs) in a CA1 pyramidal cell (DIC and Alexa Fluor 594 channel); patch and stimulating pipettes shown. Traces, two uEPSC examples (minimal stimulation) showing failures and one-quantum responses. **B**, Reversing V_h_ from -70 mv to +40 decelerates uEPSC decay: one-synapse example including a failure (5-trial average; light grey line, trace at -70 mV normalised to that at +40 mV). **C**, Summary, decay times of uEPSCs recorded at -70 mv and +40 mV, in dendrites (mean ± SEM: 1.9 ± 0.25 and 2.56 ± 0.22 ms, respectively; p < 0.003) and the soma (mean ± SEM: 3.2 ± 0.49 and 4.58 ± 0.59 ms, respectively; p < 0.002), as indicated (n = 5 cells); same colour depict recordings from the same cell. **D**, Dual-patch soma-dendrite experiment (DIC and Alexa Fluor); no detectable reverse dialysis into the dendritic pipette (dend); local sub-dendritic stimulation electrode (stim) shown. **E**, Traces, one-cell example of dendritic (top) and somatic (bottom) multi-synaptic EPSCs at -70 and + 40 mV, as indicated (10-trial average; grey line, trace at -70 mV normalised to that at +40 mV). Graph, increases in the EPSC decay time upon a switch from -70 to +40 mV (mean ± SEM: 2.68 ± 0.24 and 16.38 ±1.83 ms, respectively; n = 6, p < 0.001), recorded pairwise at dendrites and the soma, as indicated; bar, mean value; connected data point show the same cell. **F**, Traces, one-cell example of dendritic (top) and somatic (bottom) multi-synaptic EPSCs, recorded pairwise in baseline and after application of 10 µM CTZ, as indicated (10-trial average; grey line, trace in control normalised to that in CTZ). Graph, deceleration in the EPSC decay time (in ms) after CTZ application, recorded at dendrites and the soma pairwise, as indicated, (mean ± SEM: 3.13 ± 0.4 and 14.03 ±5.60 ms, respectively; n = 6, p < 0.001); other notations as **E**. **G**, One-cell example of dendritic (top) and somatic (bottom) EPSCs (10-trial average) recorded at -70 and + 40 mV, as indicated (grey line, trace at -70 mV normalised to that at +40 mV); AMPA desensitisation is blocked by 10 µM CTZ in the bath medium. **H**, Summary of experiments shown in F-G; **i**ncrease in the EPSC decay time upon a switch from -70 to +40 mV, recorded at dendrites and the soma, as indicated; connected data point depict the same recorded cell; average increases are (mean ± SEM) 6.82 ± 0.54 ms and 14.03 ± 2.29 ms (n = 6 cells; difference at p < 0.026).

Next, we set out to manipulate the uEPSC waveform, without affecting glutamate release (*I*_*h*_ and GABA signalling were blocked), using two complementary experimental designs. First, we reversed holding voltage, in soma and dendrites, from -70 mV to +40 mV (NMDA receptors were blocked by 50 µM APV). Although the kinetics of AMPA receptors is strictly voltage-independent in these cells, current reversal should retard escape of negatively charged glutamate from the synaptic cleft due to electrodiffusion, thus slowing down the AMPA receptor-mediated EPSC decay (Sylantyev *et al*., 2008; Sylantyev *et al*., 2013). Indeed, voltage reversal decelerated the decay of dendritic uEPSCs by 0.66 ± 0.09 ms (n = 5; Fig. 5B-C). This deceleration was fully consistent with the effect of glutamate electrodiffusion measured earlier in electrically compact cerebellar granule cells (Sylantyev *et al*., 2013). At the same time, voltage reversal prolonged the pairwise-recorded somatic EPSCs by 1.38 ± 0.16 ms (Fig. 5C; difference in the uEPSC decay between +40 and -70 mV at p < 0.012), which is two-fold amplification.

To extend this test to multi-synaptic activation, we increased the afferent stimulus strength while placing the stimulating pipette further away from the dendritic patch electrode (Fig. 5D). Here, the voltage asymmetry of the EPSC decay increased to 2.68 ± 0.24 ms in dendrites, and to 16.4 ± 1.8 ms in the soma (n = 6, p < 0.001; Fig. 5E). Thus, without involving *I*_*h*_ or tonic GABA current, non-linear dendritic summation can amplify small changes in the dendritic EPSC decay several-fold.

In the second experiment, we used similar settings but retarded the dendritic EPSC kinetics using cyclothiazide (CTZ), an AMPA receptor desensitisation blocker, in sub-saturation concentrations (10 µM). Again, whilst CTZ decelerated the decay of dendritic EPSCs by only 3.13 ± 0.39 ms, the slowdown at the soma was 14.0 ± 2.3 ms (n = 6, p < 0.007; Fig. 5F), which is more than a four-fold increase. Finally, we repeated the electrodiffusion experiment (Fig. 5A-E) in the presence of CTZ. As in the tests above, the depolarisation-dependent deceleration of local dendritic EPSCs (at V_m_ = +40 mV) increased more than two-fold when reaching the soma (from 6.82 ± 0.54 to 14.03 ± 2.29 ms; Fig. 5G-H).

## DISCUSSION

It has long been suggested that feedforward inhibition is a key feature enabling precise coincidence detection, and thus accurate information transfer, in central neural circuits (Pouille & Scanziani, 2001; Perez-Orive *et al*., 2002; Calixto *et al*., 2008; Pavlov *et al*., 2011). Some of these studies employed a classical experimental design in acute brain slices, in which afferent fibres are stimulated using an extracellular electrode. However, studies in the hippocampal CA3-CA1 circuit that employed either pre-post-synaptic cell pair recordings or selective optogenetic stimulation of Schaffer collaterals, documented spike-generating monophasic EPSPs in CA1 PCs, with no detectable GABAergic component (Debanne *et al*., 1996; Zhang *et al*., 2008; Kohl *et al*., 2011; Jackman *et al*., 2014). Similarly, *in vivo* recordings in hippocampal CA1 PCs appear to routinely show monophasic subthreshold EPSPs (Bahner *et al*., 2011; Kowalski *et al*., 2016). As these observations highlighted functional significance of direct excitatory inputs to CA1 PCs, it was important to understand what mechanisms can adaptively control coincidence detection of such inputs.

The combined excitatory and feed-forward inhibitory transmission in the CA3-CA1 circuit manifests itself as a prominent biphasic EPSP-IPSP response in CA1 PCs (Pouille & Scanziani, 2001; Pavlov *et al*., 2011). This response is sensitive to *I*_*h*_, which thus has a profound influence on coincidence detection in the postsynaptic pyramidal cell (Pavlov *et al*., 2011). To enable stimulation of direct excitatory inputs to CA1 PCs, here we used theta-glass bipolar electrodes that provide highly localised excitation of Schaffer collaterals. The EPSPs recorded under this protocol had no IPSP component, and upon the blockade of GABA_A_ receptors or *I*_*h*_ showed negligible changes compared to the prominent, two-fold increases in the amplitude and rise time, which has been characteristic for the case of feedforward inhibition (Pouille & Scanziani, 2001; Pavlov *et al*., 2011). These observations confirmed that our protocols enabled us to explore integration of monosynaptic inputs to CA1 PCs.

It has long been acknowledged that in CA1 PCs (and probably other principal neurons), membrane-shunting conductance carried by *I*_*h*_ plays a key role in shaping somatic response in the course of dendritic integration (Magee, 1999; Angelo *et al*., 2007; George *et al*., 2009). *I*_*h*_ has also been responsible for significant control over coincidence detection of CA3-CA1 signals in the presence of feedforward inhibition (Pavlov *et al*., 2011).

Another, well acknowledged and no less prominent source of membrane shunting, has been tonic GABAergic inhibition which depends on local [GABA] and exerts strong control over the cell spiking response (e.g., (Hausser & Clark, 1997; Semyanov *et al*., 2003; Prescott *et al*., 2006; Pavlov *et al*., 2014)). We asked therefore whether, in the absence of direct inhibitory inputs, *I*_*h*_ and tonic GABA current can regulate temporal coincidence of excitatory inputs to PCs.

Unlike the expression of *I*_*h*_, which must be a ‘stationary’ feature of individual cells, tonic GABA current depends on the dynamic equilibrium of GABA release and uptake (Glykys & Mody, 2007; Pavlov *et al*., 2014), which may vary from region to region (Lee & Maguire, 2014), reflecting local activity of neuronal networks and astroglia (Semyanov *et al*., 2004; Glykys & Mody, 2007; Woo *et al*., 2018). Indeed, we used a highly sensitive GABA sniffer patch method to demonstrate that changes in neural network activity in the slice could alter [GABA] 2-3 fold. It has previously been shown that blocking GABA transport, or indeed blocking GABA_A_ receptors, generates a comparable change of the tonic GABA current in CA1 PCs (Semyanov *et al*., 2003; Scimemi *et al*., 2005; Pavlov *et al*., 2014). We could thus employ the blockade of GABA_A_ receptors and of GABA uptake as a way to replicate activity-dependent changes in [GABA], but with the advantage of having a steady-state condition enabling coincidence detection measurements.

We found that, indeed, GABA_A_ receptor blockade and the suppression of GABA tonic current result, respectively, in the widening and the narrowing of the coincidence detection window, and that the presence of *I*_*h*_ did not seem to influence the effect of [GABA]. Among the GABA transporters, the neuronal GAT-1 type turned out to have the key contributing role. Intriguingly, expression of GABA transporters can be functionally regulated by tyrosine phosphorylation (Law *et al*., 2000), which could, in theory, provide an adaptive mechanism to regulate [GABA] and therefore coincidence detection.

Finally, we noticed that manipulations with [GABA] in our experiments could produce only a tiny (sub-millisecond or millisecond range) change in the kinetics of dendritic EPSPs. At the same time, changes in the coincidence detection window were in the range of 10-20 ms. Because passive dendritic filtering cannot explain such amplification, we employed dual soma-dendrite recordings to see whether a small change in the kinetics of local dendritic EPSCs is actively amplified at the soma, without engaging *I*_*h*_ or GABAergic signalling. To test this, we used two experimental manipulations that change the EPSC decay independently of GABA, *I*_*h*_, or glutamate release, and found significant amplification of small changes in the dendritic EPSC kinetics when the current reaches the soma. Changes in the decay of single-synapse dendritic uEPSCs were amplified approximately two-fold whereas multi-synaptic activation generated a 7-10-fold boost when recorded somatically. Thus, there appears to be an inherent mechanism of active dendritic filtering that informs the cell soma about small changes in the receptor current kinetics at dendritic synapses. Whether this mechanism plays a role in altering cell spiking behaviour depending on subtle changes in synaptic receptor composition remains an open and intriguing question.

## Acknowledgements

This study was supported by the Wellcome Trust Principal Fellowship (212251_Z_18_Z), ERC Advanced Grant (323113), and European Commission NEUROTWIN grant (857562) to DAR; University of Edinburgh Chancellor’s Fellowship to SS.

## Author contributions

SS designed and carried our electrophysiological experiments and analyses; LPS designed and carried out computer simulations; NN carried out selected physiological experiments; DAR narrated the study, designed selected experiments and simulations, carried out selected analyses, and wrote the manuscript, which was contributed by all the authors.

## Data availability statement

The raw data are available on request and will be deposited for download.

